# Cryopreservation of midbrain dopaminergic neural cells differentiated from human embryonic stem cells

**DOI:** 10.1101/2020.02.11.944272

**Authors:** Nicola J. Drummond, Karamjit Singh Dolt, Maurice A. Canham, Peter Kilbride, G. John Morris, Tilo Kunath

## Abstract

Recent advancements in protocols to differentiate human pluripotent stem cells into midbrain dopaminergic (mDA) neurons has improved the ability to model Parkinson’s disease (PD) in a dish, and has provided a scalable source of donor cells for emerging PD cell replacement therapy (CRT). However, to facilitate reproducibility, collaboration, and clinical trials it would be highly beneficial to cryopreserve committed mDA neural precursors cells in a ready-to-use format. In terms of cell manufacturing for PD CRT trials, a cryopreserved transplantation-ready mDA cell product would provide a critical opportunity for quality control, efficacy testing, and safety assessments. To address this challenge, we have compared six (6) different clinical-grade cryopreservation media and different freezing conditions for mDA neural precursor cells differentiated from two human embryonic stem cell (ESC) lines, MasterShef7 and RC17. Significant differences in cell viability were observed at 24h post-thawing, but no differences were observed immediately upon thawing. This highlights the need to check cell viability over the first 24h after thawing, and that viability of freshly thawed cells is insufficient to gauge the success of a cryopreservation protocol. Considerable apoptosis occurs in the first 24h post-thawing, and significant differences between cryopreservation procedures were only revealed during this time period. The presence of ROCK inhibitors improved cell viability at 24h for all conditions tested. A faster cooling rate of 1-2°C/min was significantly better than 0.5°C/min for all conditions tested, while rapid thawing at 37°C was not always superior to slow thawing at 4°C. Indeed, the optimal cryopreservation and thawing conditions in this study, as determined by 24h post-thaw viability, were cells frozen in PSC Cryopreservation medium at a cooling rate of 1°C/min and slow thawing at 4°C. These conditions permitted recovery of 60%-70% live cells at 24h with respect to the starting number of cryopreserved cells. Importantly, cryopreservation of mDA neural precursor cells did not alter their potential to resume differentiation into mDA neurons.

**Highlights:**

- First systematic comparison of multiple clinical-grade cryopreservation media for human ESC-derived mDA neural precursor cells
- Differences in cell viability were observed at 24h after thawing, but not immediately upon thawing
- Cooling rates of 1°C/min or 2°C/min were significantly better than 0.5°C/min for all cryopreservation conditions tested
- A slow thawing condition at 4°C was significantly better than quick thawing at 37°C for cells frozen in PSC Cryopreservation medium
- Cryopreservation of mDA cells does not significantly alter their potential to differentiate into mDA neurons

## Introduction

Parkinson’s disease (PD) is a common neurodegenerative condition characterised by progressive and selective neuronal cell loss. Although the subtypes of neurons that become dysfunctional and die in PD are diverse, the dopaminergic (DA) neurons of the *substantia nigra* are particularly affected in this condition. The embryological origin of nigral DA neurons is a population of radial glial-like cells in the floor plate of the mesencephalon [1,2]. Significant progress has been made in the last 10 years to produce floor plate cells and authentic midbrain DA (mDA) neurons from human pluripotent stem cells [3-6]. The ability of mDA neurons differentiated from human embryonic stem cells (ESCs) and induced pluripotent stem cells (iPSCs) to rescue *in vivo* pre-clinical models of PD has been extensively investigated [4,6,7]. Human ESC-derived mDA cells were demonstrated to be functionally equivalent to human fetal ventral midbrain tissue in the rat 6-hydroxydopamine (6-OHDA) lesion model of PD [8]. Furthermore, human iPSC-derived mDA neural precursor cells, FACS-sorted for the floor plate marker CORIN, could rescue a macaque model of PD established by 1-methyl-4-phenyl-1,2,3,6-tetrahydropyridine (MPTP) lesion [7]. These international efforts are culminating in the first cell replacement therapy (CRT) clinical trials for PD [9]. The improved mDA differentiation protocols have also enhanced the ability to model PD in a dish. Single-cell RNAseq of human ESC/iPSC-derived mDA neurons generated by the floor plate protocol showed significant overlap with multiple human fetal mDA cell types [10].

Differentiation of human ESCs/iPSCs into mDA neurons is a complex and multi-stage process, and it is known that different iPSC lines from the same patient can have significantly different propensities to produce mDA neurons [11]. Furthermore, the positional identity of floor plate cells produced from human ESCs/iPSCs is highly sensitive to small changes in WNT signalling [6]. A cryopreserved mDA neural precursor cell population could provide a quality-controlled cell bank from which mDA neuronal differentiation and maturation can be conducted. This will reduce variability across experiments, and facilitate collaborations with multiple laboratories. Furthermore, a cryopreserved transplantation-ready mDA cell product will provide a time window for safety testing and assessment of efficacy for CRT clinical trials. A frozen cell product would also provide a practical solution for multi-centre trials, and allow direct comparison of results across sites and indeed patients.

Cryopreservation of primary rat fetal mesencephalic tissue resulted in a greater than 50% loss of viability compared to non-frozen cells, but the surviving neurons, when grafted into the rat 6-OHDA lesion model, were able to ameliorate the amphetamine-induced rotation phenotype [12]. However, attempts to cryopreserve human fetal mesencephalic tissue prior to grafting were less successful with more than 90% loss of viable mDA cells compared to non-frozen controls, and no significant rescue of amphetamine-induced rotations [13]. More recently, successful cryopreservation of human ESC/iPSC-derived mDA cells using a floor plate protocol have been reported [14,15]. Furthermore, commercial cryopreserved human iPSC-derived mDA cells (iCell DopaNeurons) have been thawed and directly transplanted into rat and non-human primate lesion models of PD, and rescue of amphetamine-induced rotations was observed in the rat model, and survival and maturation into DA neurons was observed in the monkey model [16]. Here we build upon these findings and present the first systematic report comparing multiple clinical-grade cryopreservation media and cryopreservation conditions for human ESC-derived mDA neural precursor cells. We propose that high-quality cryopreserved human mDA cells will improve reproducibility across experiments, facilitate collaborations, and provide a critical step to advance CRT for Parkinson’s disease.

## Results

Two clinical-grade human ESC lines, MasterShef7 (MShef7) and RC17, were differentiated into mDA neural precursor cells and then into neurons using a modified 2D floor plate protocol [4,17]. Briefly, human ESCs were plated onto a matrix of Laminin-111 in the presence of dual-Smad inhibition (SB431542 + LDN193189), 0.9-1 μM GSK3β inhibitor (CHIR9902), and a high concentration of recombinant Sonic hedgehog (SHH, 600 ng/ml). FGF8b and heparin were added at day 9 of differentiation, and brain-derived growth factor (BDNF) and glial-derived growth factor (GDNF) were added at day 11 (Figure 1A). The differentiating mDA cells can be conveniently cryopreserved at day 11 or day 16 of differentiation, since these are time-points when the cells are normally lifted and re-plated in the protocol. However, for all experiments in this study cells were frozen at day 16 since this the stage they can be transplanted into pre-clinical rat models of Parkinson’s [6], and cells at this point of mDA commitment are ideal for clinical translation. In order to examine the *in vitro* differentiation potential of the cryopreserved cells, the mDA precursors are thawed onto Laminin-111 in the presence of neurotrophic and neuronal maturation factors to resume differentiation into mDA neurons up to day 45 (Figure 1A). At the point of freezing (day 16), the vast majority of cells expressed the mDA markers, LMX1A and FOXA2 (Figure 1B), and greater than 85% of the cells were positive for the ventral floor plate marker, CORIN, by flow cytometry (Figure 1C).

**Figure 1.**
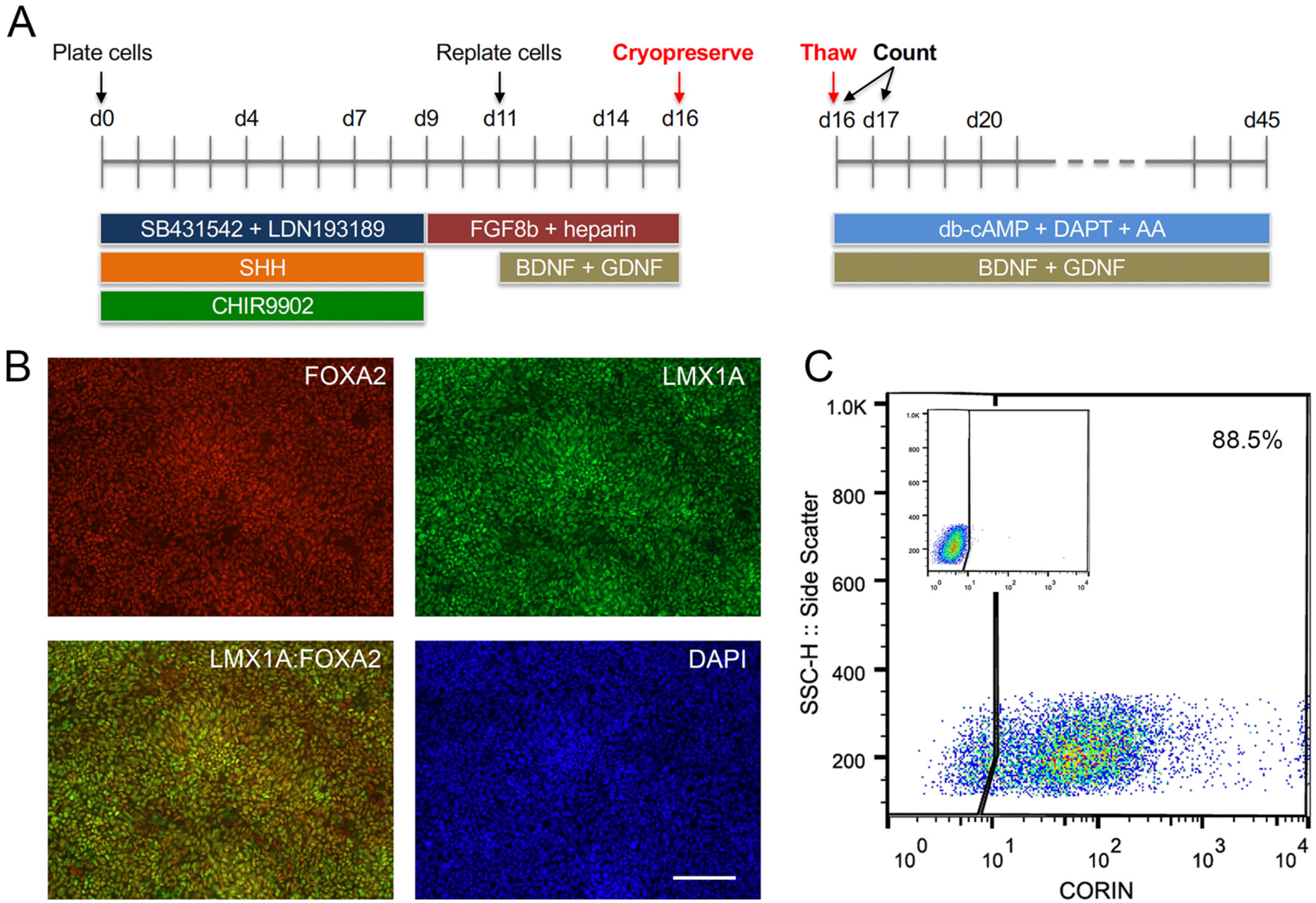
Differentiation of human embryonic stem cells (ESCs) into midbrain dopaminergic (mDA) precursors for cryopreservation. **(A)** Schematic of the mDA differentiation protocol. Human ESCs are plated as clumps on day 0 in the presence of dual SMAD inhibitors, SB431542 + LDN193189, Sonic hedgehog (SHH), and the GSK3β inhibitor, CHIR9902. At day 9 the medium is switched to contain only FGF8b and the co-factor heparin, and cells are lifted and re-plated at day 11 when the neurotrophic factors BDNF and GDNF are added. At day 16 the cells are lifted and counted for cryopreservation. Live/dead cell counts were performed immediately upon thawing and at 24h after thawing (day 17). mDA neuronal maturation was conducted in the presence of BDNF, GDNF, db-cAMP, ascorbic acid (AA), and the Notch inhibitor DAPT, up to day 45. **(B)** Immunostaining for midbrain floor plate markers, FOXA2 and LMX1A, at day 16 revealed near homogenous expression at the point of cryopreservation. Scale bar, 120 μm. **(C)** FACS analysis for CORIN expression at day 16 showed the majority of cells expressed this ventral floor plate marker. The control FACS plot without primary antibody is shown in the inset.

Human ESC-derived mDA cells were cryopreserved at a density of ∼1×10^7^ cells/ml in six (6) different clinical-grade cryopreservation media (Table 1). The media investigated include (i) STEM-CELLBANKER^®^ (SCB), (ii) Synth-a-Freeze™ Medium (SYF), (iii) PSC Cryopreservation Medium (PSC), (iv) CryoStor^®^ CS10 Freeze Medium, (v) CryoStor^®^ CS5 Freeze Medium, and (vi) Cellvation Cryopreservation Medium (CV). CS5 contained 5% DMSO (v/v), while CV is a DMSO-free cryopreservation medium. The other 4 cryopreservation media contained 10% DMSO (v/v). HypoThermosol^®^, a 4°C clinical-grade hibernation medium, was used as a negative control for these experiments. Cooling rates were varied from 0.5°C/min to 2°C/min using a VIA Freeze controlled-rate freezer starting from 4°C and stopping at -80°C before transferring vials to liquid nitrogen storage (Figure 2). The mDA precursor cells were then either thawed slowly in air at 4°C or rapidly in a water-bath set to 37°C prior to plating in mDA neuronal differentiation conditions. Cell viability was assessed by Trypan blue exclusion immediately upon thawing, and at 24h post-thawing, since cryopreservation-induced apoptosis occurs 12h-24h after thawing [18]. In order to gain a full account of the fate of all cells, both the floating cells and attached cells were counted at 24h post-thawing (Figure 2).

**Table 1.**
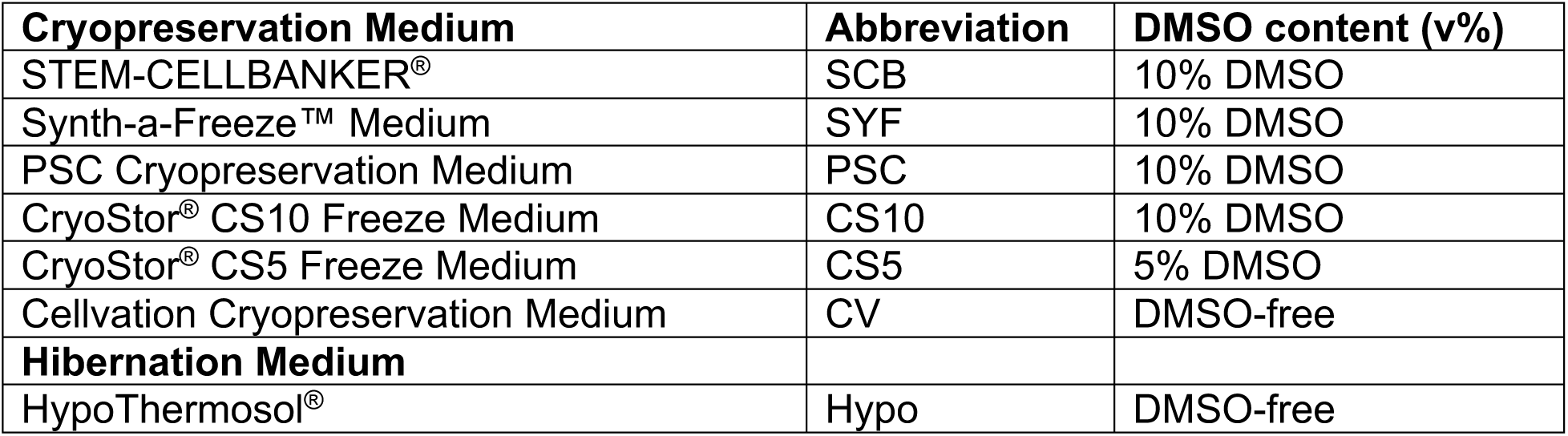
List of media used for cryopreservation.

| **Cryopreservation Medium** | **Abbreviation** | **DMSO content (v%)** |
| --- | --- | --- |
| STEM-CELLBANKER® | SCB | 10% DMSO |
| Synth-a-Freeze™ Medium | SYF | 10% DMSO |
| PSC Cryopreservation Medium | PSC | 10% DMSO |
| CryoStor® CS10 Freeze Medium | CS10 | 10% DMSO |
| CryoStor® CS5 Freeze Medium | CS5 | 5% DMSO |
| Cellvation Cryopreservation Medium | CV | DMSO-free |
| **Hibernation Medium** |  |  |
| HypoThermosol® | Hypo | DMSO-free |

**Figure 2.**
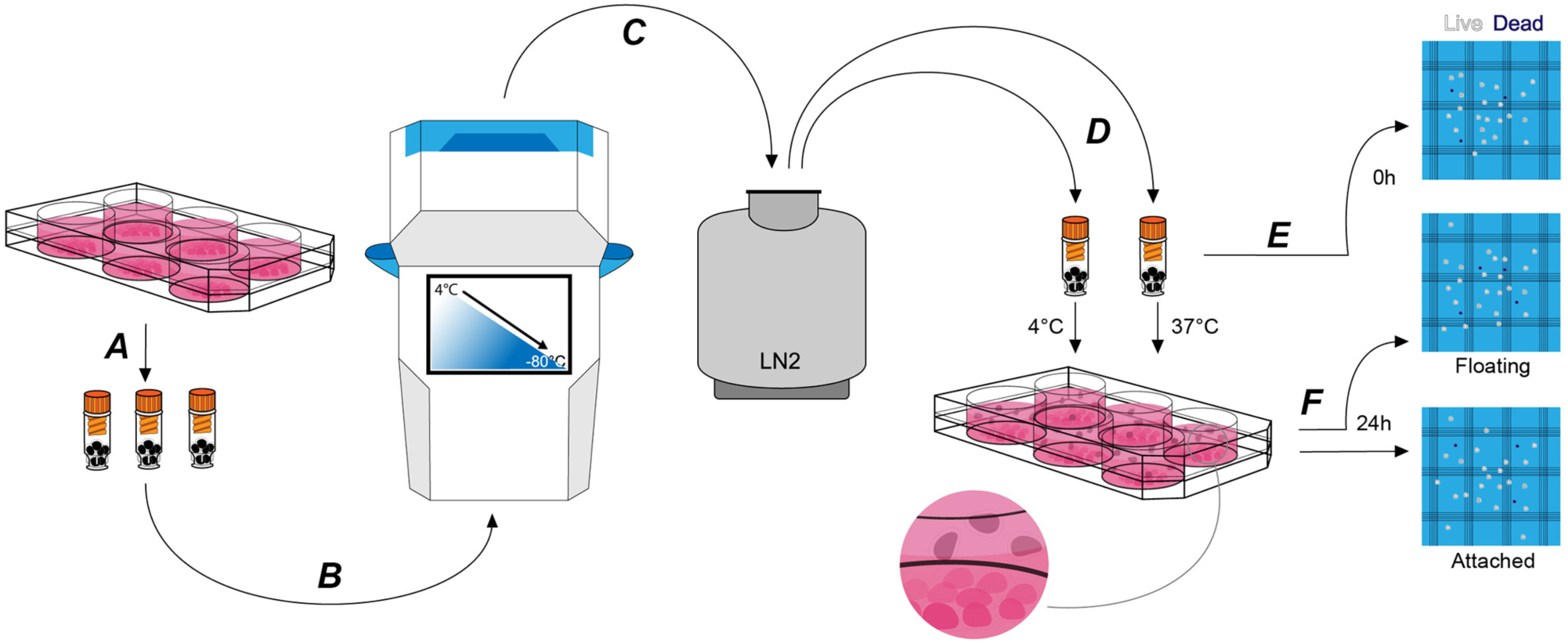
Cryopreservation experimental procedure. **(A)** At day 16 of mDA differentiation, cells were lifted, counted, and resuspended in a clinical-grade cryopreservation medium prior to transferring to FluidX cryovials. **(B)** Cryovials were placed in the VIA Freeze controlled-rate freezer and cells were cryopreserved at a constant rate of cooling between 0.5°C/min to 2°C/min to a final temperature of -80°C. **(C)** Cryovials were transferred to the vapour phase of liquid nitrogen for long term storage. **(D)** Cryopreserved cells were thawed at 4°C or 37°C. **(E)** A live/dead cell count was performed immediately after thawing (0h count) prior to plating the cells. **(F)** 24h after plating cells (day 17 of differentiation) the non-adherent floating cells and the adherent attached cells were subjected to live/dead cell counts (“24h – Floating” count and “24h – Attached” count, respectively).

Rho-associated kinase (ROCK) inhibitors have been shown to significantly improve cell survival of single cells upon passaging and after cryopreservation [19]. We first compared the commercial formulation, RevitaCell, which contains a ROCK inhibitor and antioxidants, with Y27632 (10 μM), a cell-permeable ROCK inhibitor. At 24h post-thawing both RevitaCell and Y27632 significantly increased the number of attached live mDA cells (Figure 3A), and significantly reduced the number of floating cells (Figure 3B). There were no significant differences between RevitaCell and Y27632 by these measures. Since RevitaCell is manufactured at cGMP-grade for medical devices (21 CFR Part 820 and ISO 13485), it was used for all subsequent experiments.

**Figure 3.**
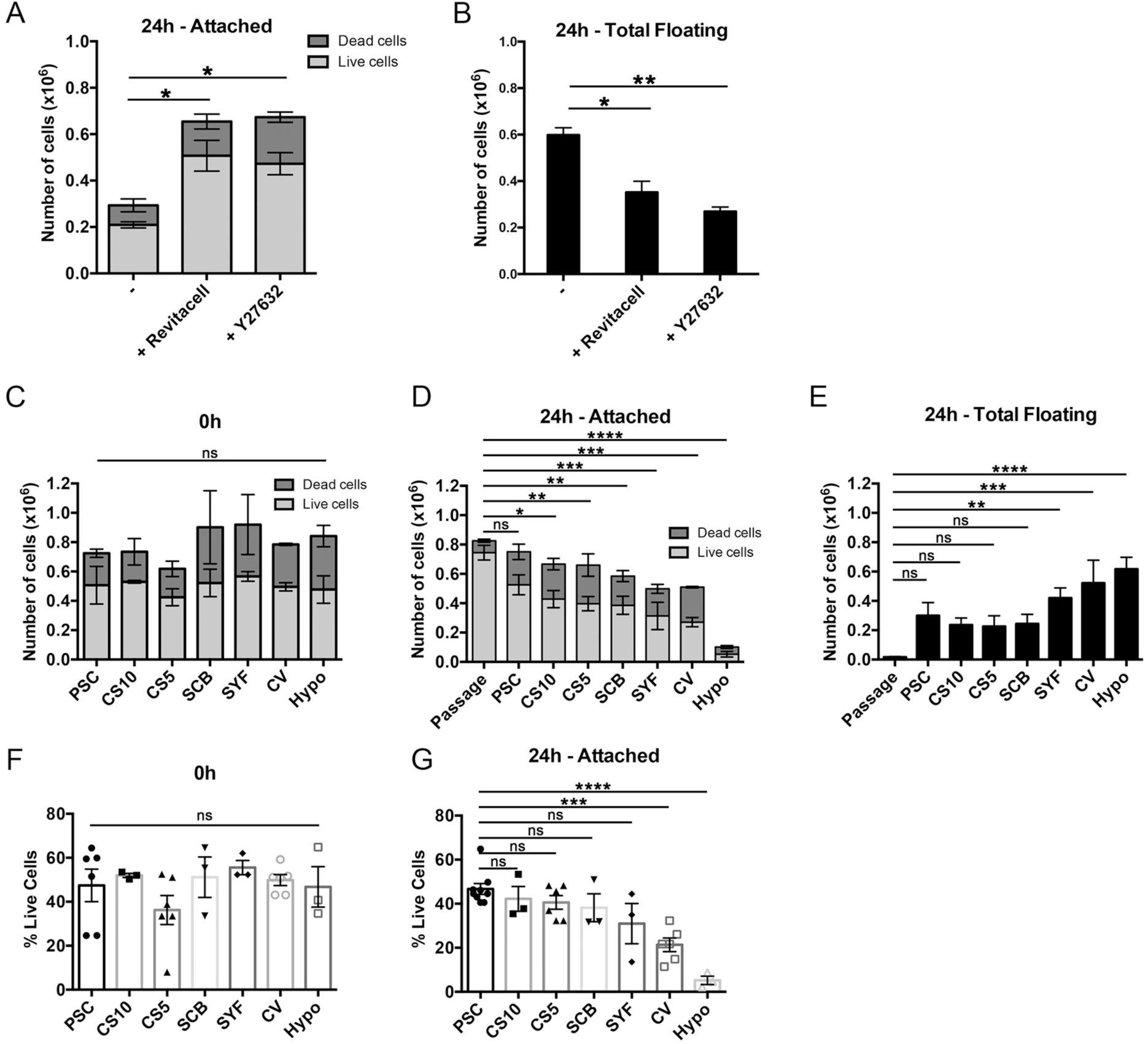
Assessment of cell viability after cryopreservation in the presence or absence of ROCK inhibitors, and using 7 different clinical-grade media. **(A**,**B)** Cell viability of mDA cells after cryopreservation in the absence of ROCK inhibitors or in the presence of RevitaCell™ or Y27632. (A) Live/dead *<u>attached</u>* cell numbers at 24h after thawing, and (B) total number of *<u>floating</u>* cells at 24h (n=2). **(C)** Comparison of cell viability after cryopreservation in PSC Cryopreservation Medium (PSC), CryoStor^®^ CS10 Freeze Media (CS10), CryoStor^®^ CS5 Freeze Media (CS5), STEM-CELLBANKER^®^ (SCB), Synth-a-Freeze™ Medium (SYF), Cellvation Cryopreservation Medium (CV), and Hypothermosol^®^ (Hypo) immediately after thawing. **(D**,**E)** At 24h post-thawing live/dead cell counts of the attached cells (D) and total floating cells (E) were determined (n=3), and compared to freshly passaged cells (n=6). **(F**,**G)** Live cells as a percentage of initial cell number frozen immediately upon thawing (F), and attached cells at 24h post-thawing (G). ns, not significant, * p < 0.05, ** p < 0.01, *** p < 0.001, **** p < 0.0001 one-way ANOVA with Tukey’s multiple comparisons test.

We conducted a side-by-side comparison of 6 commercially-available clinical-grade cryopreservation media, and a clinical-grade DMSO-free hibernation medium, HypoThermasol^®^ (Hypo) (Table 1). Cell viability immediately upon thawing was not significantly different for all cryopreservation media, nor was it significantly different to the hibernation medium (Figure 3C). However, at 24h post-thawing there were significant differences in cell viability for different cryopreservation media, and the hibernation medium, Hypo, gave the lowest level of cell viability and the most floating cells (Figure 3D,E). The best performing medium, PSC Cryopreservation Medium, was the only one that was not significantly different from freshly passaged cells (Figure 3D). Cellvation, a DMSO-free cryopreservation medium, was not significantly worse that the DMSO-containing cryopreservation media, while Hypo was significantly worse than PSC, CS10, CS5, and SCB cryopreservation media (Supplementary Table S2). The number of floating cells at 24h post-thawing was significantly higher for cells frozen in SYF, CV, and Hypo media, but not for cells cryopreserved in PSC, CS10, CS5, or SCB media (Figure 3E). When the data is analyzed as a percentage of the initial number of frozen cells, there are no significance differences in the percent of live cells immediately upon thawing for any of the media (Figure 3F), but at 24h post-thawing the DMSO-free media, CV and Hypo, have significantly lower percentages of live cells than the best-performing medium, PSC (Figure 3G). While the performance of CV and Hypo were not significantly different from each other by this measure, Hypo was significantly poorer than all the DMSO-containing cryopreservation media, while the DMSO-free CV media was not statistically different from SCB and SYF media, which contain 10% DMSO (v/v) (Supplementary Table S2).

We next investigated three cooling rates and two thawing conditions for the best-performing medium, PSC, as well as for CS5 and CV media, which contained 5% DMSO and 0% DMSO, respectively. For PSC media, 1°C/min and 2°C/min were significantly better than 0.5°C/min at 24h post-thawing, while there were no significant differences for cells frozen in CS5 medium (Figure 4A,B). For the DMSO-free CV medium, the fastest freezing rate of 2°C/min was significantly better than the slowest rate of 0.5°C/min (Figure 4C). Although it is common practice to thaw cells rapidly in a warm water-bath, we found that slow thawing in cool conditions (10 min at 4°C) was significantly better than fast thawing at 37°C for cells frozen in PSC media (Figure 4D). However, mDA cells cryopreserved in CS5 or CV media did not show significant differences between the cool and warm thawing conditions (Figure 4E,F). Neural mDA precursor cells differentiated from another clinical-grade human ESC line, RC17, also exhibited excellent survival 24h post-thawing after cryopreservation in PSC medium (Figure 4G). In agreement with the MShef7 data, slow thawing at 4°C was equivalent, if not superior, to rapid thawing at 37°C for RC17-derived mDA cells frozen in PSC medium (Figure 4G). Cells frozen in the 5% DMSO medium, CS5, did not recover as well as cells cryopreserved in PSC medium, however, there was a trend towards better survival in the fast 37°C thawing condition for mDA cells differentiated from both MShef7 and RC17 cell lines (Figure 4E,H).

**Figure 4.**
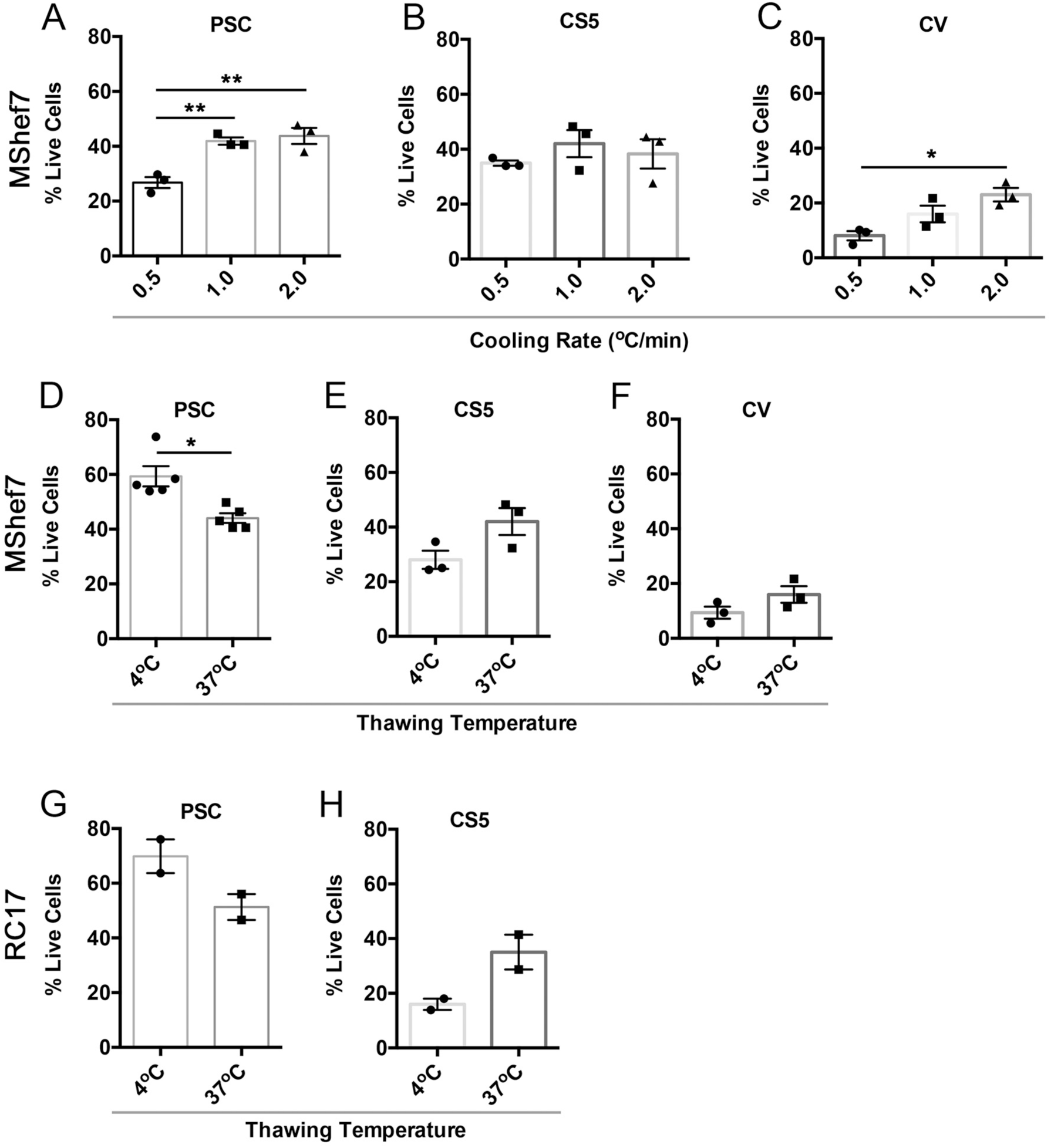
Cell recovery at different cooling rates and thawing conditions. **(A-C)** Percentage of live attached cells recovered 24h after thawing for cells cryopreserved at different cooling rates, 0.5, 1.0, and 2.0°C per minute, in PSC (A), CS5 (B), or CV (C) cryopreservation media. * p < 0.05, ** p < 0.01, one-way ANOVA with Tukey’s multiple comparisons test. **(D-F)** Cell recovery at 24h post-thawing for mDA cells frozen in PSC (n=5), CS5 (n=3), or CV (n=3) cryopreservation media and thawed at 4°C or 37°C. Cells were originally * p < 0.05, unpaired *t*-test with Welch’s correction. **(G**,**H)** Cell recovery of RC17-derived mDA cells at different thaw rates (37°C vs 4°C) in PSC and CS5 cryopreservation medium (n=2).

Next we directly compared the capacity of non-frozen to frozen mDA cells to produce dopaminergic neurons up to 45 days of differentiation. There were no gross differences in the production of neurons for cells frozen at day 16 of differentiation when compared to non-frozen cells in terms of morphology or expression of the pan-neuronal marker βIII-tubulin and the dopaminergic marker, tyrosine hydroxylase (TH) (Figure 5A). Gene expression analysis of *TH*, and two mDA markers, *NURR1* and *PITX3*, also did not reveal any significant differences between frozen and non-frozen mDA cells differentiated from both MShef7 and RC17 human ESC lines (Figure 5B).

**Figure 5.**
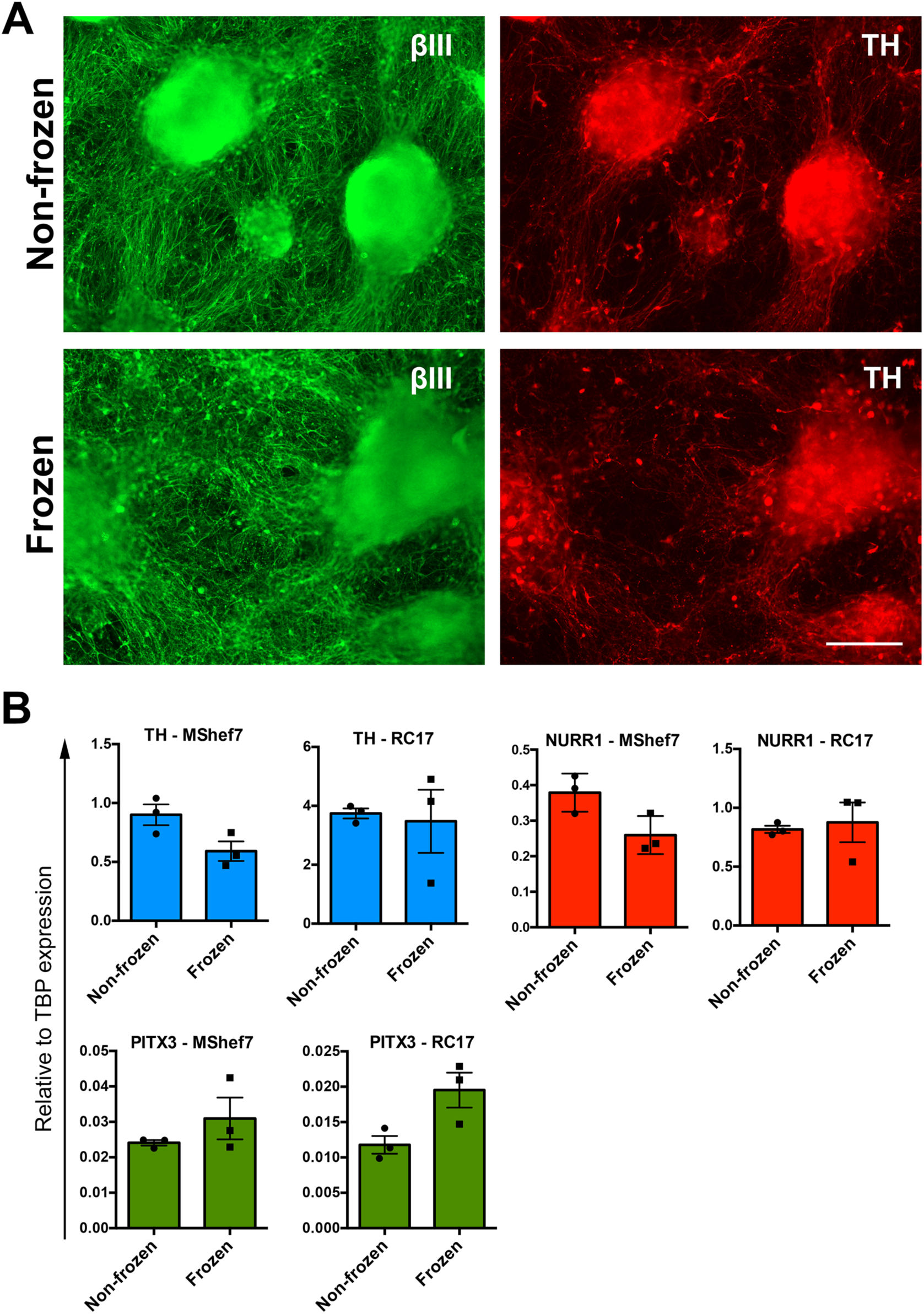
Marker analysis of continuously differentiated mDA neurons versus neurons that were cryopreserved at day 16 as mDA neural precursor cells. **(A)** Immunofluorescence staining for the pan-neuronal marker, βIII tubulin (βIII), and dopaminergic marker, tyrosine hydroxylase (TH), at day 45 of differentiation with or without freezing at day 16. Scale bar, 220 μm. **(B)** RT-qPCR gene expression analysis of mDA markers *TH, NURR1*, and *PITX3* at day 42 of differentiation from two human ESC lines, MShef7 and RC17 (n=3).

## Discussion

Human pluripotent stem cells are unique because they can produce cell types of all three germ layers in culture, maintain high chromosomal stability, and are practically immortal [20,21]. These properties make human ESCs and iPSCs ideal for modelling disease in a dish, as well as providing an unlimited source of cells for regenerative medicine applications [22,23]. Numerous protocols exist to differentiate human ESCs/iPSCs into a diverse range of cell types, including many ectodermal and neuronal subtypes [24]. Recent advances in differentiation of ventral mDA neurons has provided a significant opportunity to model PD in a dish, and it has accelerated the route to cell replacement therapies for this condition [9]. Although current differentiation protocols can produce high yields of mDA neurons, cell line-to-cell line differences continue to exist, as well as other sources of variation due in part to the complexity of the protocol [25]. It is well described that different human ESC and iPSC lines have variable potency of directed differentiation into neurons and other cell types [26-28]. One solution to this is to produce a quality-controlled cryopreserved bank of mDA neural progenitors from which all experiments or transplantations are initiated from. This is a similar concept to the CryoPause methodology where banks of human ESCs or iPSCs are cryopreserved at one passage in a ready-to-differentiate format, without the need for expansion [29]. Here we propose that neural progenitors, frozen in a ready-to-use format, will significantly improve the reproducibility of disease modelling, as well as facilitate efforts towards cell replacement therapy. The feasibility of a cryopreserved ‘paused’ differentiated cell product was demonstrated when frozen iCell DopaNeurons were shown to be capable of rescuing the 6-OHDA lesion rat model of PD [16].

Since there is significant cell loss associated with freezing and thawing of cells, we took a systematic approach to examine multiple aspects of the cryopreservation process in order to maximise the yield of viable mDA cells. To this end we identified six clinical-grade cryopreservation media to test against each other (Table 1), and investigated a clinical-grade 4°C hibernation medium, HypoThermosol^®^ (Hypo), which has been used to cryopreserve cells in the literature [30]. We investigated different cooling rates with a VIA Freeze™ Duo controlled-rate freezer, and compared rapid versus slow thawing of cells (Figure 2). We focused our cryopreservation experiments on day 16 of the mDA protocol, since this is the day that we lift the differentiating cells for re-plating, and this is the optimal stage of maturity for transplantation into the rat 6-OHDA lesion model of PD [6]. We first investigated the addition of a selective Rho-associated coiled-coil kinase (ROCK) inhibitor, Y27632, and the supplement Revitacell that contains a ROCK inhibitor, to the recovery medium. We found that both agents significantly improved cell survival in the first 24h after thawing (Figure 3A,B). These observations are in agreement with published work demonstrating that inhibition of ROCK in the first 12h after passaging or thawing of cells prevents apoptosis in a number of cell types including ESCs and neurons [19,31].

Surprisingly, there were no significant differences in cell viability immediately after thawing cells for any of the cryopreservation media, including the hibernation medium, Hypo (Figure 3C). Differences only emerged when cell viability was quantified at 24h post-thawing (Figure 3D). In this case Hypo was the worst-performing cryopreservation media, followed by SYF and CV, while the best-performing media, PSC, was not significantly different to freshly passaged cells (Figure 3D). The observations of cell viability at 0h and 24h post-thawing are in agreement with data describing cryopreservation-induced delayed onset of cell death [18,30,32]. Quantification of apoptosis in the first 48h after thawing MDCK cells identified the highest rates of apoptosis to be between 12h-24h [18]. This work and our observations strongly suggest that measuring cell viability at 24h post-thawing is more informative than cell viability measurements immediately upon thawing. Furthermore, an indirect measure of the efficiency of a cryopreservation protocol can been inferred from the number of floating cells at 24h post-thawing (Figure 3E). This identified Hypo, CV, and SYF as significantly poorer than other cryopreservation media for mDA cells. Although DMSO-free CV medium was the worst-performing cryopreservation medium tested, it was consistently better than Hypo by all measures at 24h post-thawing. DMSO is a highly effective cryopreservation agent, but other solutes, such as glycerol, can provide a measure of cryoprotection [33].

Varying the cooling rate resulted in significant differences in 24h post-thaw cell viability for PSC and CV cryopreservation media, where faster cooling rates (1°C/min and 2°C/min) were generally better than a slow cooling rate of 0.5°C/min (Figure 4A,C). Varying the cooling rate for cells frozen in CS5 media did not result in any significant differences in cell viability (Figure 4B). Slow cooling rates are known to avoid intracellular ice crystal formation in part by allowing the cells sufficient time to lose water [34], however excessively slow cooling may result in DMSO toxicity. Optimal cooling rates are dependent on cell type, but also cryoprotectant as we have seen here. Some cryoprotectants containing extracellular solutes will enhance cell dehydration, therefore allowing sufficient dehydration at more rapid cooling rates. Other solutes may be more toxic so more sensitive to slower cooling rates where cells will remain in the cryoprotectant longer before reaching long term storage conditions [35].

We next investigated rapid versus slow thawing conditions. Cryopreserved cells are typically thawed quickly in a 37°C water-bath, and slow thawing is usually considered undesirable, which is supported by limited experimental data from the 1970s [36,37]. However, warming rates for cryopreserved mammalian somatic cells was recently re-visited, and no impact of thawing conditions could be identified for cells cooled slowly at a rate of 1°C/min [38]. Only when cells were frozen quickly, 10°C/min or higher, was rapid thawing beneficial. Such rapid rates of cooling result in only partial ice crystal formation, and rapid thawing some damage due to re-crystallization on warming. If cells are cryopreserved slowly in optimal conditions, then the warming rate had little impact of viability as ice crystals fully develop during the cooling [38]. In agreement with this data, we did not observe a significant improvement in cell viability for cells thawing quickly in a 37°C water-bath compared to cells thawed slowly in air at 4°C. In fact, slow thawing for MShef7-derived mDA cells was significantly better than rapid thawing (Figure 4D). An additional reason why slow thawing might be beneficial could be the non-uniform nature of thawing frozen liquids, which could be exaggerated during rapid thawing. Ice does not melt uniformly at the microscale, and it can melt in some places, and re-freeze in other places, leading to ice re-crystallization that can damage cells. Slow thawing could reduce these micro-fluctuations. Recent studies with HepG2 cells suggested slow thawing up to -10°C, and then rapid thawing from -10°C to 4°C is beneficial due to the reduction of micro-fluctuations [39]. Finally, rapid thawing may not be beneficial due to the potential for increased toxicity of DMSO, especially if temperatures approach 37°C before wash-out [40]. An ideal thawing protocol for some cell types may consist of slow thawing to 4°C, followed by immediate dilution of cells in cold medium to reduce the toxic effects of DMSO prior to centrifugation.

We next directly compared the differentiation potential of cryopreserved cells to cells that were not frozen up to 45 days of mDA differentiation. Although our analysis was not comprehensive, there were no gross morphological differences between the mDA neurons produced from frozen versus non-frozen cells, and we did not observed any significant differences in the expression of dopaminergic markers, *TH, NURR1*, and *PITX3* for two different human ESC lines, RC17 and MShef7 (Figure 5).

This study provides the first systematic investigation of cryopreservation conditions for human mDA progenitor cells. We found that assessment of cell viability at 24h post-thawing, and not immediately upon thawing, was essential to discriminant between good and poor cryopreservation conditions. Furthermore, we challenge the notion that rapid thawing of cells is necessarily better than slow thawing conditions. Indeed, non-linear cooling and warming conditions could further improve the cell viability of human mDA cells, and warrants further investigation. Other parameters that can be tested moving forward include cell density, total freezing volume, and cryovial type (material, wall thickness), which were all kept constant in this study. High-quality cryopreserved mDA cells with high post-thaw viability will be a valuable resource for the neuroscience research community, and will greatly facilitate the efforts for a cell replacement therapy for Parkinson’s.

## Materials & Methods

### Human embryonic stem cell culture

Approval for the use of MasterShef7 (MShef7) and RC17 human embryonic stem cells (ESCs) was granted by the MRC Steering Committee for the UK Stem Cell Bank and for the Use of Stem Cell Lines (ref. SCSC13-18 for MShef7 and ref. SCSC13-19 for RC17). Human ESCs were maintained in StemMACS™ iPS-Brew XF (iPS-B, Miltenyi Biotec) on Laminin-521 (L521, 5 μg/ml, Biolamina) coated plates. Once 70-90% confluent they were passaged as clumps using EDTA (0.5 mM, Thermo Fisher Scientific).

### Midbrain dopaminergic differentiation

Self-renewing human ESCs were lifted with EDTA (0.5 mM), counted and plated for differentiation at 40,000 cells/cm^2^ on Laminin-111-coated plates (L111, 5 μg/ml, Biolamina) in 50% Neurobasal™ medium (Thermo Fisher Scientific), 50% DMEM/F12 (Thermo Fisher Scientific), B27 (1:50, Thermo Fisher Scientific), N2 (1:100, Thermo Fisher Scientific) and 2 mM L-glutamine (Thermo Fisher Scientific) with Sonic hedgehog (Shh-C24II, 600 ng/ml, R&D), CHIR99021 (0.9 μM or 1 μM, Miltenyi Biotec), SB431542 (10 μM, Tocris), LDN193189 (100 nM, Stemgent) and Y27632 (10 μM, Tocris). The culture medium was changed on day 2 with the above medium without Y27632. On day 4 and day 7 medium consisted of 50% Neurobasal, 50% DMEM/F12, B27 (1:100) and N2 (1:200), 2 mM L-glutamine supplemented with Sonic hedgehog (600 ng/ml), CHIR99021 (0.9 μM or 1 μM), SB431542 (10 μM), LDN193189 (100 nM). On day 9 medium was supplemented with FGF8b (100 ng/ml, R&D) and heparin (1 μg/ml, Sigma). On day 11 cells were lifted using Accutase (Sigma) and replated at 800,000 cells/cm^2^ on L111-coated plates in Neurobasal medium supplemented with B27 (1:50), 2 mM L-glutamine, BDNF (20 ng/ml, Peprotech), GDNF (10 ng/ml, Peprotech), ascorbic acid (0.2 mM, Sigma), FGF8b (100 ng/ml), heparin (1 μg/ml), and Y27632 (10 μM). Cells were fed on day 14 with day 11 medium without Y27632. On day 16 cells were lifted with Accutase and either (i) cryopreserved or (ii) replated at 800,000 cells/cm^2^ on L111-coated plates in Neurobasal medium with B27 (1:50) and 2 mM L-glutamine supplemented with BDNF (20 ng/ml), GDNF (10 ng/ml), ascorbic acid (0.2 mM), dibutyryl cyclic AMP (db-cAMP, 0.5 mM, Sigma) and DAPT (1 μM, Tocris). From day 18 onwards medium was changed every 2-3 days.

### Cryopreservation

At day 16 of differentiation cells were lifted using Accutase and counted using TC20™ Automated Cell counter (BioRad). Cells were centrifuged (300g, 3 min) and resuspended in 100 μl of the appropriate freezing medium (Table 1) in 0.5-ml tubes (FluidX) at 1.02×10^6^ cells per cryovial. These were moved to the VIA Freeze™ Duo (Asymptote, GE Healthcare) and were frozen using the following protocol: 4°C hold for 10 minutes and then temperature reduced at 0.5°C per minute, 1°C per minute or 2°C per minute until - 80°C was reached. Cryovials were transported on dry ice and transferred to the vapour phase liquid nitrogen for long term storage.

### Thawing and cell counting

Cells were thawed by exposing the vial to 37°C water for 1-2 minutes or by placing the vial in air at 4°C for approximately 10 minutes. The cell suspension was removed from the cryovial and placed onto 1 ml Neurobasal™ medium with B27 (1:50) and 2 mM L-glutamine and centrifuged (300g, 3 min). The supernatant was removed and the cell pellet was resuspended in 300 μl of day 16 differentiation medium supplemented with either Y27632 (10 μM) or RevitaCell™ (1X, Thermo Fisher Scientific). An aliquot of cells was taken for cell counting (0h cell count) prior to plating in L111-coated 48-well plates (Corning). 24 hours later the conditioned medium was collected, and combined with a DPBS wash of the cells to collect all the floating cells. This was centrifuged (300g, 3 min) and cells were resuspended in a small volume (20 μl -120 μl) of Neurobasal medium with B27 (1:50), 2 mM L-glutamine and counted in the presence of Trypan Blue using the TC20™ Automated Cell counter (Bio-Rad) (24h floating cell count). The adherent cells were lifted using Accutase and similarly counted (24h attached cell count).

### Immunofluorescence staining

Cells were fixed with 4% formaldehyde for 20 minutes and washed three times in PBS. Immunocytochemistry was performed by addition of blocking buffer (0.1% Triton X-100, 2% goat serum or donkey serum in PBS) for 30 minutes. Primary antibodies TH (1:1000, rabbit, Millipore), TuJ1 (1:1000, mouse IgG2a, R&D), LMX1A (1:2000, rabbit, Millipore), and FOXA2 (1:100, goat, Santa Cruz) were incubated with fixed cells overnight at 4°C and then washed three times in PBS with 0.1% Triton X-100. Secondary antibodies (Thermo Fisher Scientific) in blocking buffer were incubated with cells for 2 hours in the dark at room temperature. They were washed three times in PBS with Triton X-100 and incubated with DAPI (10 μg/ml, Thermo Fisher Scientific) before imaging on an Olympus IX51 inverted microscope.

### FACS

Cells were lifted using Accutase and diluted with Neurobasal medium with B27 (1:50) and 2 mM L-glutamine. Cells were centrifuged (2150g, 1.5 min), and resuspended in FACS Buffer (PBS + 2% FBS) with CORIN antibody (1:200, rat, R&D), and incubated for 15 minutes on ice. The primary antibody was omitted for the control FACS. Cells were then centrifuged and washed in FACS Buffer and incubated on ice (15 min) with secondary antibody donkey anti-rat IgG Alexa Fluor-488 (Thermo Fisher Scientific). Cells were centrifuged and washed in FACS Buffer and flow cytometry data was collected using the FACS Calibur (BD Biosciences) and post-acquisition analysis using FlowJo software.

### RT-qPCR

RNA extraction was performed using the MasterPure™ Complete DNA and RNA Purification Kit (Epicentre, MC85200), according to manufacturer’s instructions. RNA concentration was quantified using a NanoDrop spectrophotometer. Total RNA (1 μg) was used for cDNA synthesis. RNase-free water was added to the RNA to give a 10 μl sample. The samples were incubated with 1 μl dNTP mix (10 mM, Thermo Fisher Scientific) and 1 μl Random Primer Mix (60 μM, NEB) at 65°C for 5 minutes and then chilled on ice. After a brief centrifugation, 4 μl 5x First-strand buffer (Thermo Fisher Scientific), 2 μl 0.1 M DTT (Thermo Fisher Scientific) and 1 μl RNaseOUT (40 units/μl, Thermo Fisher Scientific) were added. The contents were incubated at 37°C for 2 minutes. Then 1 μl M-MLV reverse transcriptase (200 units/μl, Thermo Fisher Scientific) or 1 μl RNase-free water for the negative reverse transcriptase sample was added. This was mixed and incubated at room temperature for 10 minutes when it was moved to 37°C for 1 hour. The reaction was inactivated by incubation at 90°C for 10 minutes. The cDNA mix was placed on ice and 80 μl RNase-free water was added. qPCR was performed using the Roche LightCycler® 480 System with the Universal Probe Library (UPL) (Roche). The Roche UPL Assay design centre was used to design intron-spanning primers with a specific UPL probe for each gene (*TBP* F-gaacatcatggatcagaacaaca R-atagggattccgggagtcat, Probe 87; *NURR1* F-atttcctcgaaaacgcctgt R-catactgcgcctgaacacaa, Probe 41; *PITX3* F-tgtcagacgctggcactc R-ccgaggccttttctgagtc, Probe 24; *TH* F-gattccccgtgtggagtaca R-aagcaaaggcctccaggt, Probe 12). Reactions (10 μl) containing primers, UPL Probe, LightCycler® 480 Probes Master mix (Roche) and PCR water were performed in 384-well plates as described in the manufacturer’s instructions. The results were normalized to RNA levels of TATA-binding protein (*TBP*).

## Acknowledgements

This work was funded by MRC RMRC grant (MR/K017276/1) and MRC Confidence-in-concept awards to TK and The Cure Parkinson’s Trust.

## Data accessibility

All raw data for graphs in Figures 3, 4, and 5 can obtained from LabArchives.com at https://doi.org/10.25833/8a4w-4y22.

## Authors’ contribution

TK designed the study and wrote the paper. NJD performed experiments and data analysis. KSD and MAC provided reagents and optimised protocols. PK and GJM contributed to study design, data analysis, and loan of VIA Freeze™ Dou controlled-rate freezer. All authors edited and approved the manuscript.

## References

[1] Ono Y, Nakatani T, Sakamoto Y, Mizuhara E, Minaki Y, Kumai M, Hamaguchi A, Nishimura M, Inoue Y, Hayashi H, Takahashi J, Imai T (2007) Differences in neurogenic potential in floor plate cells along an anteroposterior location: midbrain dopaminergic neurons originate from mesencephalic floor plate cells. Development 134, 3213–3225.

[2] Bonilla S, Hall AC, Pinto L, Attardo A, Götz M, Huttner WB, Arenas E (2008) Identification of midbrain floor plate radial glia-like cells as dopaminergic progenitors. Glia 56, 809–820.

[3] Fasano CA, Chambers SM, Lee G, Tomishima MJ, Studer L (2010) Efficient Derivation of Functional Floor Plate Tissue from Human Embryonic Stem Cells. Cell Stem Cell 6, 336–347.

[4] Kriks S, Shim J-W, Piao J, Ganat YM, Wakeman DR, Xie Z, Carrillo-Reid L, Auyeung G, Antonacci C, Buch A, Yang L, Beal MF, Surmeier DJ, Kordower JH, Tabar V, Studer L (2011) Dopamine neurons derived from human ES cells efficiently engraft in animal models of Parkinson’s disease. Nature 480, 547–551.

[5] Xi J, Liu Y, Liu H, Chen H, Emborg ME, Zhang S-C (2012) Specification of midbrain dopamine neurons from primate pluripotent stem cells. Stem Cells 30, 1655–1663.

[6] Kirkeby A, Grealish S, Wolf DA, Nelander J, Wood J, Lundblad M, Lindvall O, Parmar M (2012) Generation of regionally specified neural progenitors and functional neurons from human embryonic stem cells under defined conditions. Cell Rep 1, 703–714.

[7] Kikuchi T, Morizane A, Doi D, Magotani H, Onoe H, Hayashi T, Mizuma H, Takara S, Takahashi R, Inoue H, Morita S, Yamamoto M, Okita K, Nakagawa M, Parmar M, Takahashi J (2017) Human iPS cell-derived dopaminergic neurons function in a primate Parkinson’s disease model. Nature 548, 592–596.

[8] Grealish S, Diguet E, Kirkeby A, Mattsson B, Heuer A, Bramoulle Y, Van Camp N, Perrier AL, Hantraye P, Björklund A, Parmar M (2014) Human ESC-derived dopamine neurons show similar preclinical efficacy and potency to fetal neurons when grafted in a rat model of Parkinson’s disease. Cell Stem Cell 15, 653–665.

[9] Barker RA, Parmar M, Studer L, Takahashi J (2017) Human Trials of Stem Cell-Derived Dopamine Neurons for Parkinson’s Disease: Dawn of a New Era. Cell Stem Cell 21, 569–573.

[10] La Manno G, Gyllborg D, Codeluppi S, Nishimura K, Saltó C, Zeisel A, Borm LE, Stott SRW, Toledo EM, Villaescusa JC, Lönnerberg P, Ryge J, Barker RA, Arenas E, Linnarsson S (2016) Molecular Diversity of Midbrain Development in Mouse, Human, and Stem Cells. Cell 167, 566–580.e19.

[11] Devine MJ, Ryten M, Vodicka P, Thomson AJ, Burdon T, Houlden H, Cavaleri F, Nagano M, Drummond NJ, Taanman J-W, Schapira AH, Gwinn K, Hardy J, Lewis PA, Kunath T (2011) Parkinson’s disease induced pluripotent stem cells with triplication of the α-synuclein locus. Nat Commun 2, 440.

[12] Sauer H, Frodl EM, Kupsch A, Bruggencate ten G, Oertel WH (1992) Cryopreservation, survival and function of intrastriatal fetal mesencephalic grafts in a rat model of Parkinson’s disease. Exp Brain Res 90, 54–62.

[13] Frodl EM, Duan WM, Sauer H, Kupsch A, Brundin P (1994) Human embryonic dopamine neurons xenografted to the rat: effects of cryopreservation and varying regional source of donor cells on transplant survival, morphology and function. Brain Res. 647, 286–298.

[14] Niclis JC, Gantner CW, Alsanie WF, McDougall SJ, Bye CR, Elefanty AG, Stanley EG, Haynes JM, Pouton CW, Thompson LH, Parish CL (2017) Efficiently Specified Ventral Midbrain Dopamine Neurons from Human Pluripotent Stem Cells Under Xeno-Free Conditions Restore Motor Deficits in Parkinsonian Rodents. Stem Cells Transl Med 6, 937–948.

[15] Leitner D, Ramamoorthy M, Dejosez M, Zwaka TP (2019) Immature mDA neurons ameliorate motor deficits in a 6-OHDA Parkinson’s disease mouse model and are functional after cryopreservation. Stem Cell Res 41, 101617.

[16] Wakeman DR, Hiller BM, Marmion DJ, McMahon CW, Corbett GT, Mangan KP, Ma J, Little LE, Xie Z, Perez-Rosello T, Guzman JN, Surmeier DJ, Kordower JH (2017) Cryopreservation Maintains Functionality of Human iPSC Dopamine Neurons and Rescues Parkinsonian Phenotypes In Vivo. Stem Cell Reports 9, 149–161.

[17] Kirkeby A, Nolbrant S, Tiklova K, Heuer A, Kee N, Cardoso T, Ottosson DR, Lelos MJ, Rifes P, Dunnett SB, Grealish S, Perlmann T, Parmar M (2017) Predictive Markers Guide Differentiation to Improve Graft Outcome in Clinical Translation of hESC-Based Therapy for Parkinson’s Disease. Cell Stem Cell 20, 135–148.

[18] Baust JM, Vogel MJ, Van Buskirk R, Baust JG (2001) A molecular basis of cryopreservation failure and its modulation to improve cell survival. Cell Transplant 10, 561–571.

[19] Watanabe K, Ueno M, Kamiya D, Nishiyama A, Matsumura M, Wataya T, Takahashi JB, Nishikawa S, Nishikawa S-I, Muguruma K, Sasai Y (2007) A ROCK inhibitor permits survival of dissociated human embryonic stem cells. Nat. Biotechnol. 25, 681–686.

[20] Thomson JA, Itskovitz-Eldor J, Shapiro SS, Waknitz MA, Swiergiel JJ, Marshall VS, Jones JM (1998) Embryonic stem cell lines derived from human blastocysts. Science 282, 1145–1147.

[21] Takahashi K, Tanabe K, Ohnuki M, Narita M, Ichisaka T, Tomoda K, Yamanaka S (2007) Induction of pluripotent stem cells from adult human fibroblasts by defined factors. Cell 131, 861–872.

[22] Cohen DE, Melton D (2011) Turning straw into gold: directing cell fate for regenerative medicine. Nat. Rev. Genet. 12, 243–252.

[23] Tabar V, Studer L (2014) Pluripotent stem cells in regenerative medicine: challenges and recent progress. Nat. Rev. Genet. 15, 82–92.

[24] Tchieu J, Zimmer B, Fattahi F, Amin S, Zeltner N, Chen S, Studer L (2017) A Modular Platform for Differentiation of Human PSCs into All Major Ectodermal Lineages. Cell Stem Cell 21, 399–410.e7.

[25] Nolbrant S, Heuer A, Parmar M, Kirkeby A (2017) Generation of high-purity human ventral midbrain dopaminergic progenitors for in vitro maturation and intracerebral transplantation. Nat Protoc 12, 1962–1979.

[26] Hu B-Y, Weick JP, Yu J, Ma L-X, Zhang X-Q, Thomson JA, Zhang S-C (2010) Neural differentiation of human induced pluripotent stem cells follows developmental principles but with variable potency. Proceedings of the National Academy of Sciences 107, 4335–4340.

[27] Osafune K, Caron L, Borowiak M, Martinez RJ, Fitz-Gerald CS, Sato Y, Cowan CA, Chien KR, Melton DA (2008) Marked differences in differentiation propensity among human embryonic stem cell lines. Nat. Biotechnol. 26, 313–315.

[28] Wu H, Xu J, Pang ZP, Ge W, Kim KJ, Blanchi B, Chen C, Südhof TC, Sun YE (2007) Integrative genomic and functional analyses reveal neuronal subtype differentiation bias in human embryonic stem cell lines. Proc. Natl. Acad. Sci. U.S.A. 104, 13821–13826.

[29] Wong KG, Ryan SD, Ramnarine K, Rosen SA, Mann SE, Kulick A, De Stanchina E, Muller F-J, Kacmarczyk TJ, Zhang C, Betel D, Tomishima MJ (2017) CryoPause: A New Method to Immediately Initiate Experiments after Cryopreservation of Pluripotent Stem Cells. Stem Cell Reports 9, 355–365.

[30] Baust JM, Van Buskirk, Baust JG (2000) Cell viability improves following inhibition of cryopreservation-induced apoptosis. In Vitro Cell. Dev. Biol. Anim. 36, 262–270.

[31] Lingor P, Tönges L, Pieper N, Bermel C, Barski E, Planchamp V, Bähr M (2008) ROCK inhibition and CNTF interact on intrinsic signalling pathways and differentially regulate survival and regeneration in retinal ganglion cells. Brain 131, 250–263.

[32] Baust JM, Snyder KK, VanBuskirk RG, Baust JG (2009) Changing paradigms in biopreservation. Biopreserv Biobank 7, 3–12.

[33] Lovelock JE, Bishop MW (1959) Prevention of freezing damage to living cells by dimethyl sulphoxide. Nature 183, 1394–1395.

[34] Mazur P (1963) Kinetics of water loss from cells at subzero temperatures and the likelihood of intracellular freezing. J. Gen. Physiol. 47, 347–369.

[35] Fuller BJ (2004) Cryoprotectants: the essential antifreezes to protect life in the frozen state. Cryo Letters 25, 375–388.

[36] Harris LW, Griffiths JB (1977) Relative effects of cooling and warming rates on mammalian cells during the freeze-thaw cycle. Cryobiology 14, 662–669.

[37] Akhtar T, Pegg DE, Foreman J (1979) The effect of cooling and warming rates on the survival of cryopreserved L-cells. Cryobiology 16, 424–429.

[38] Baboo J, Kilbride P, Delahaye M, Milne S, Fonseca F, Blanco M, Meneghel J, Nancekievill A, Gaddum N, Morris GJ (2019) The Impact of Varying Cooling and Thawing Rates on the Quality of Cryopreserved Human Peripheral Blood T Cells. Sci Rep 9, 3417–13.

[39] Kilbride P, Lamb S, Gibbons S, Bundy J, Erro E, Selden C, Fuller B, Morris J (2017) Cryopreservation and re-culture of a 2.3 litre biomass for use in a bioartificial liver device. PLoS ONE 12, e0183385.

[40] Morris TJ, Picken A, Sharp DMC, Slater NKH, Hewitt CJ, Coopman K (2016) The effect of Me2SO overexposure during cryopreservation on HOS TE85 and hMSC viability, growth and quality. Cryobiology 73, 367–375.

