## Supplementary Information for "Cryopreservation of midbrain dopaminergic neural cells differentiated from human embryonic stem cells"

**Supplementary Table S1.** Reagents and catalogue numbers

| Reagent | Company | Cat. No. |
|---|---|---|
| DPBS with calcium, magnesium | Sigma | 14040-083 |
| Cell culture plates | Corning | 3516, 3524, 3548, 3595 |
| StemMACS iPS-Brew XF | Miltenyi Biotec | 130-107-086 |
| UltraPure 0.5 M EDTA | Thermo Fisher Scientific | 15575-038 |
| Accutase | Sigma | A6964 |
| Laminin-521 | BioLamina | LN521 |
| Laminin-111 | BioLamina | LN111 |
| DMEM/F12, no glutamine | Thermo Fisher Scientific | 21331-020 |
| Neurobasal Medium | Thermo Fisher Scientific | 21103-049 |
| B27 supplement | Thermo Fisher Scientific | 12587-010 |
| N2 supplement | Thermo Fisher Scientific | 17502-048 |
| L-Glutamine | Thermo Fisher Scientific | 25030123 |
| Y27632 | Tocris | 1254 |
| SB431542 | Millipore | 616461 |
| LDN193189 | Miltenyi Biotec | 130-103-925 |
| Shh-C24II | R&D Systems | 1845-SH-500 |
| CHIR99021 | Miltenyi Biotec | 130-103-926 |
| FGF8b | R&D Systems | 423-F8/CF |
| heparin | Sigma | H3149 |
| ascorbic acid | Sigma | A4403 |
| brain-derived neurotrophic factor (BDNF) | Peprotech | 450-02 |
| glial cell line-derived neurotrophic factor (GDNF) | Peprotech | 450-10 |

| | | |
|---|---|---|
| dibutyryl cyclic AMP (db-cAMP) | Sigma | D0627 |
| DAPT | Tocris | 2634 |
| formaldehyde solution 37-41% | Fisher | F/1501/PB08 |
| donkey serum | Sigma | D9663 |
| goat serum | Sigma | G9023 |
| Triton-X 100 | Fisher | BP-151-100 |
| PBS | Thermo Fisher Scientific | 18912-014 |
| rabbit anti-tyrosine hydroxylase | Millipore | AB152 |
| goat anti-HNF3β/FOXA2 (M20) | Santa Cruz | sc-6554 |
| rabbit anti-LMX1A | Millipore | ab10533 |
| mouse anti-β-III tubulin | R&D Systems | MAB1195 |
| rat anti-CORIN | R&D Systems | MAB2209 |
| donkey anti-rabbit IgG Alexa Fluor-488 | Thermo Fisher Scientific | A21206 |
| donkey anti-goat IgG Alexa Fluor-568 | Thermo Fisher Scientific | A11057 |
| donkey anti-rat IgG Alexa Fluor-488 | Thermo Fisher Scientific | A21208 |
| goat anti-mouse IgG2a Alexa Fluor-488 | Thermo Fisher Scientific | A21131 |
| goat anti-rabbit Alexa Fluor-555 | Thermo Fisher Scientific | A21428 |
| DAPI | Thermo Fisher Scientific | D1306 |
| Fetal Bovine Serum (FBS) | Thermo Fisher Scientific | 10270-106 |
| MasterPure™ Complete DNA and RNA Purification Kit | Epicentre | MC85200 |
| M-MLV Reverse Transcriptase | Thermo Fisher Scientific | 28025013 |
| Nuclease-Free Water | Thermo Fisher Scientific | AM9937 |
| dNTP Set | Thermo Fisher Scientific | 10297018 |
| Random Primer Mix | New England Biolabs | S1330S |
| 0.1 M DTT | Thermo Fisher Scientific | 707265ML |
| RNaseOUT | Thermo Fisher Scientific | 10777019 |

| | | |
|---|---|---|
| LightCycler® 480 Probes Master mix | Roche | 04707494001 |
| LightCycler® 480 Multiwell Plate 384 | Roche | 4729749001 |
| 0.5ml EasyTrack 2D-coded screw top tubes | FluidX | 66-52325-Z6 |
| CryoStor® CS10 Freeze Media | BioLife Solutions | 210102 |
| CryoStor® CS5 Freeze Media | BioLife Solutions | 205102 |
| HypoThermosol® | BioLife Solutions | 101373 |
| PSC Cryopreservation Kit | Thermo Fisher Scientific | A2644601 |
| RevitaCell ™ Supplement (100x) | Thermo Fisher Scientific | A2644501 |
| Cellvation Cryopreservation Medium | MP Biomedicals | SKU 0920300M2 |
| CTS Synth-a-Freeze™ medium | Thermo Fisher Scientific | A1371301 |
| STEM-CELLBANKER | Amsbio (Zenoaq) | 11890 |
| Cell Counting Slides for TC10™/TC20™ Cell Counter | Bio-Rad | 1450015 |
| Trypan blue | Sigma | T8154 |

**Supplementary Table S2.** Statistical tests

**For Figure 3A:** 24h Attached live cell counts

**Test:** one-way ANOVA with Tukey’s multiple comparisons

| Comparison | Adjusted P value | Significance |
|---|---|---|
| - vs. Revitacell | 0.0298 | * |
| - vs. Y27632 | 0.0415 | * |
| RevitaCell vs. Y27632 | ns | |

**For Figure 3B:** 24h Floating total cell counts

**Test:** one-way ANOVA with Tukey’s multiple comparisons

| Comparison | Adjusted P value | Significance |
|---|---|---|
| - vs. Revitacell | 0.0119 | * |
| - vs. Y27632 | 0.0052 | ** |
| RevitaCell vs. Y27632 | ns | |

**For Figure 3C:** 0h Live cell counts

**Test:** one-way ANOVA with Tukey’s multiple comparisons

| Comparison | Adjusted P value | Significance |
|---|---|---|
| PSC vs. CS10 | ns | |
| PSC vs. CS5 | ns | |
| PSC vs. SCB | ns | |
| PSC vs. SYF | ns | |
| PSC vs. CV | ns | |
| PSC vs. Hypo | ns | |
| CS10 vs. CS5 | ns | |
| CS10 vs. SCB | ns | |
| CS10 vs. SYF | ns | |
| CS10 vs. CV | ns | |
| CS10 vs. Hypo | ns | |
| CS5 vs. SCB | ns | |
| CS5 vs. SYF | ns | |
| CS5 vs. CV | ns | |
| CS5 vs. Hypo | ns | |
| SCB vs. SYF | ns | |

| | |
|---|---|
| SCB vs. CV | ns |
| SCB vs. Hypo | ns |
| SYF vs. CV | ns |
| SYF vs. Hypo | ns |
| CV vs. Hypo | ns |

**For Figure 3D:** 24h Attached live cell counts

**Test:** one-way ANOVA with Tukey's multiple comparisons

| **Comparison** | **Adjusted P value** | **Significance** |
|---|---|---|
| Passage vs. PSC | ns | |
| Passage vs. CS10 | 0.0101 | * |
| Passage vs. CS5 | 0.0042 | ** |
| Passage vs. SCB | 0.0031 | ** |
| Passage vs. SYF | 0.0004 | *** |
| Passage vs. CV | 0.0001 | *** |
| Passage vs. Hypo | < 0.0001 | **** |
| PSC vs. CS10 | ns | |
| PSC vs. CS5 | ns | |
| PSC vs. SCB | ns | |
| PSC vs. SYF | ns | |
| PSC vs. CV | ns | |
| PSC vs. Hypo | 0.0007 | *** |
| CS10 vs. CS5 | ns | |
| CS10 vs. SCB | ns | |
| CS10 vs. SYF | ns | |
| CS10 vs. CV | ns | |
| CS10 vs. Hypo | 0.0078 | ** |
| CS5 vs. SCB | ns | |
| CS5 vs. SYF | ns | |
| CS5 vs. CV | ns | |
| CS5 vs. Hypo | 0.0167 | * |
| SCB vs. SYF | ns | |
| SCB vs. CV | ns | |
| SCB vs. Hypo | 0.0216 | * |

| | |
|---|---|
| SYF vs. CV | ns |
| SYF vs. Hypo | ns |
| CV vs. Hypo | ns |

**For Figure 3E:** 24h Floating total cell counts

**Test:** one-way ANOVA with Tukey's multiple comparisons

| **Comparison** | **Adjusted P value** | **Significance** |
|---|---|---|
| Passage vs. PSC | ns | |
| Passage vs. CS10 | ns | |
| Passage vs. CS5 | ns | |
| Passage vs. SCB | ns | |
| Passage vs. SYF | 0.0076 | ** |
| Passage vs. CV | 0.0007 | *** |
| Passage vs. Hypo | < 0.0001 | **** |
| PSC vs. CS10 | ns | |
| PSC vs. CS5 | ns | |
| PSC vs. SCB | ns | |
| PSC vs. SYF | ns | |
| PSC vs. CV | ns | |
| PSC vs. Hypo | ns | |
| CS10 vs. CS5 | ns | |
| CS10 vs. SCB | ns | |
| CS10 vs. SYF | ns | |
| CS10 vs. CV | ns | |
| CS10 vs. Hypo | 0.0384 | * |
| CS5 vs. SCB | ns | |
| CS5 vs. SYF | ns | |
| CS5 vs. CV | ns | |
| CS5 vs. Hypo | 0.0317 | * |
| SCB vs. SYF | ns | |
| SCB vs. CV | ns | |
| SCB vs. Hypo | 0.0448 | * |
| SYF vs. CV | ns | |
| SYF vs. Hypo | ns | |

| | |
|---|---|
| CV vs. Hypo | ns |

**For Figure 3G:** 24h Percent live cells

**Test:** one-way ANOVA with Tukey's multiple comparisons

| Comparison | Adjusted P value | Significance |
|---|---|---|
| PSC vs. CS10 | ns | |
| PSC vs. CS5 | ns | |
| PSC vs. SCB | ns | |
| PSC vs. SYF | ns | |
| PSC vs. CV | 0.0002 | *** |
| PSC vs. Hypo | < 0.0001 | **** |
| CS10 vs. CS5 | ns | |
| CS10 vs. SCB | ns | |
| CS10 vs. SYF | ns | |
| CS10 vs. CV | 0.0318 | * |
| CS10 vs. Hypo | 0.0003 | *** |
| CS5 vs. SCB | ns | |
| CS5 vs. SYF | ns | |
| CS5 vs. CV | 0.0112 | * |
| CS5 vs. Hypo | < 0.0001 | **** |
| SCB vs. SYF | ns | |
| SCB vs. CV | ns | |
| SCB vs. Hypo | 0.0015 | ** |
| SYF vs. CV | ns | |
| SYF vs. Hypo | 0.0183 | * |
| CV vs. Hypo | ns | |

**For Figure 4A:** 24h Percent live cells for different cooling rates in PSC

**Test:** one-way ANOVA with Tukey's multiple comparisons

| Comparison | Adjusted P value | Significance |
|---|---|---|
| 0.5°C/min vs. 1°C/min | 0.0067 | ** |
| 0.5°C/min vs. 2°C/min | 0.0038 | ** |
| 1°C/min vs. 2°C/min | ns | |

**For Figure 4C:** 24h Percent live cells for different cooling rates in CV

**Test:** one-way ANOVA with Tukey's multiple comparisons

| Comparison | Adjusted P value | Significance |
|---|---|---|
| 0.5°C/min vs. 1°C/min | ns | |
| 0.5°C/min vs. 2°C/min | 0.0119 | * |
| 1°C/min vs. 2°C/min | ns | |

**For Figure 4D:** 24h Percent live cells for different thawing conditions in PSC

**Test:** unpaired *t*-test with Welch's correction

| Comparison | Adjusted P value | Significance |
|---|---|---|
| 4°C vs. 37°C | 0.0067 | * |
